## Supplementary figures and images for "The nanoCUT&RUN technique visualizes telomeric chromatin in Drosophila"

### supplemental figure S1

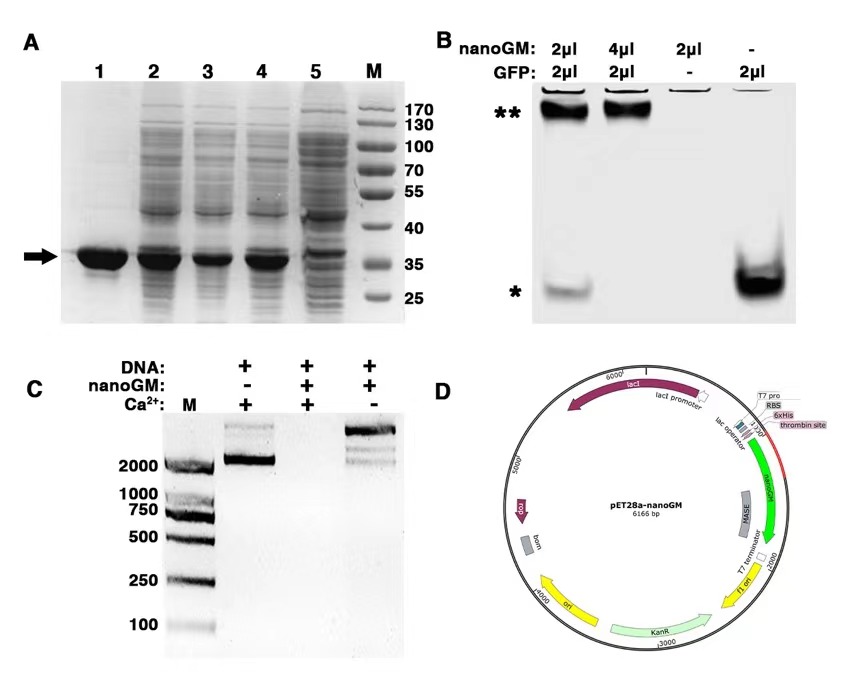

### supplemental figure S2

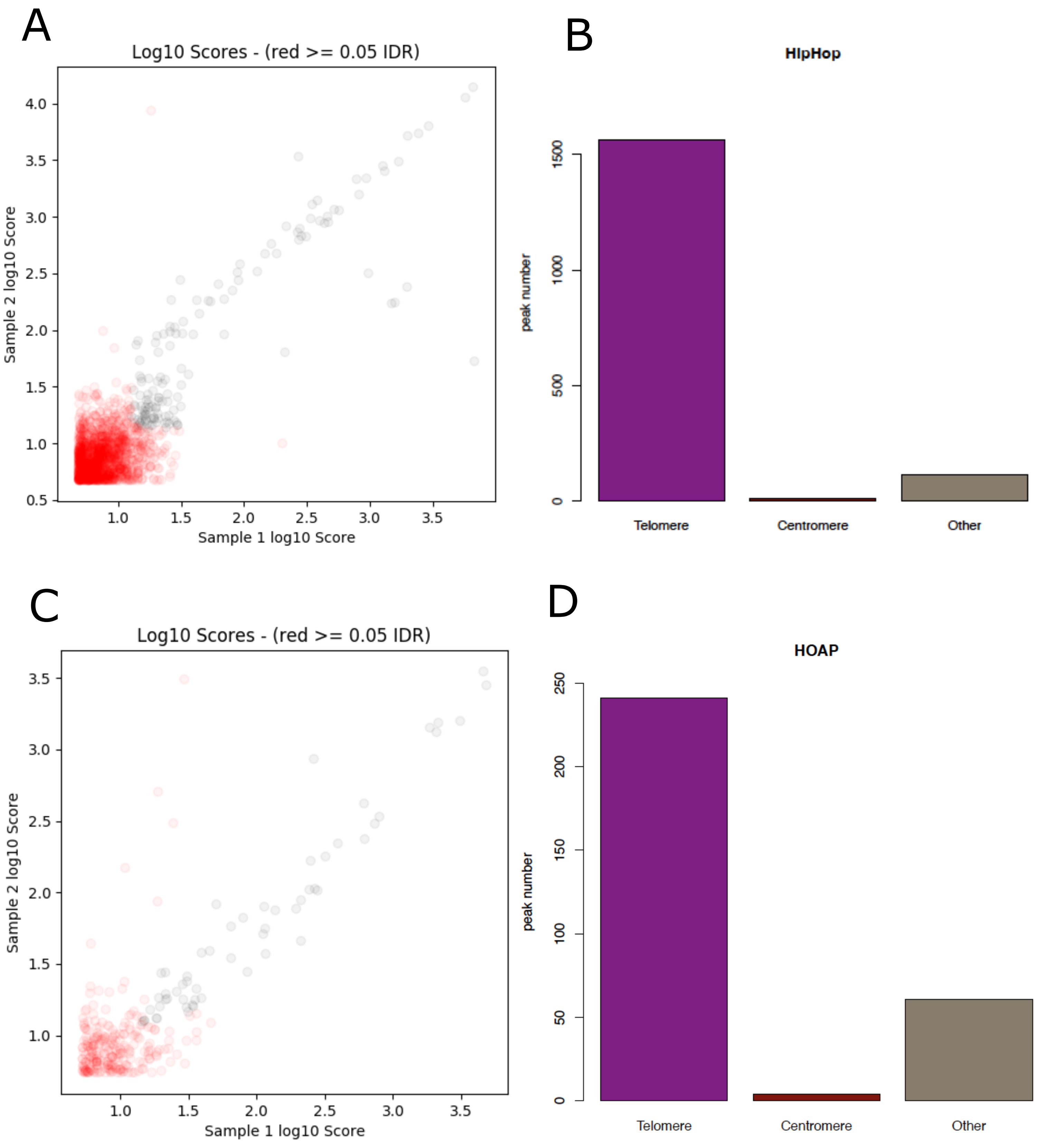

### supplemental figure S3

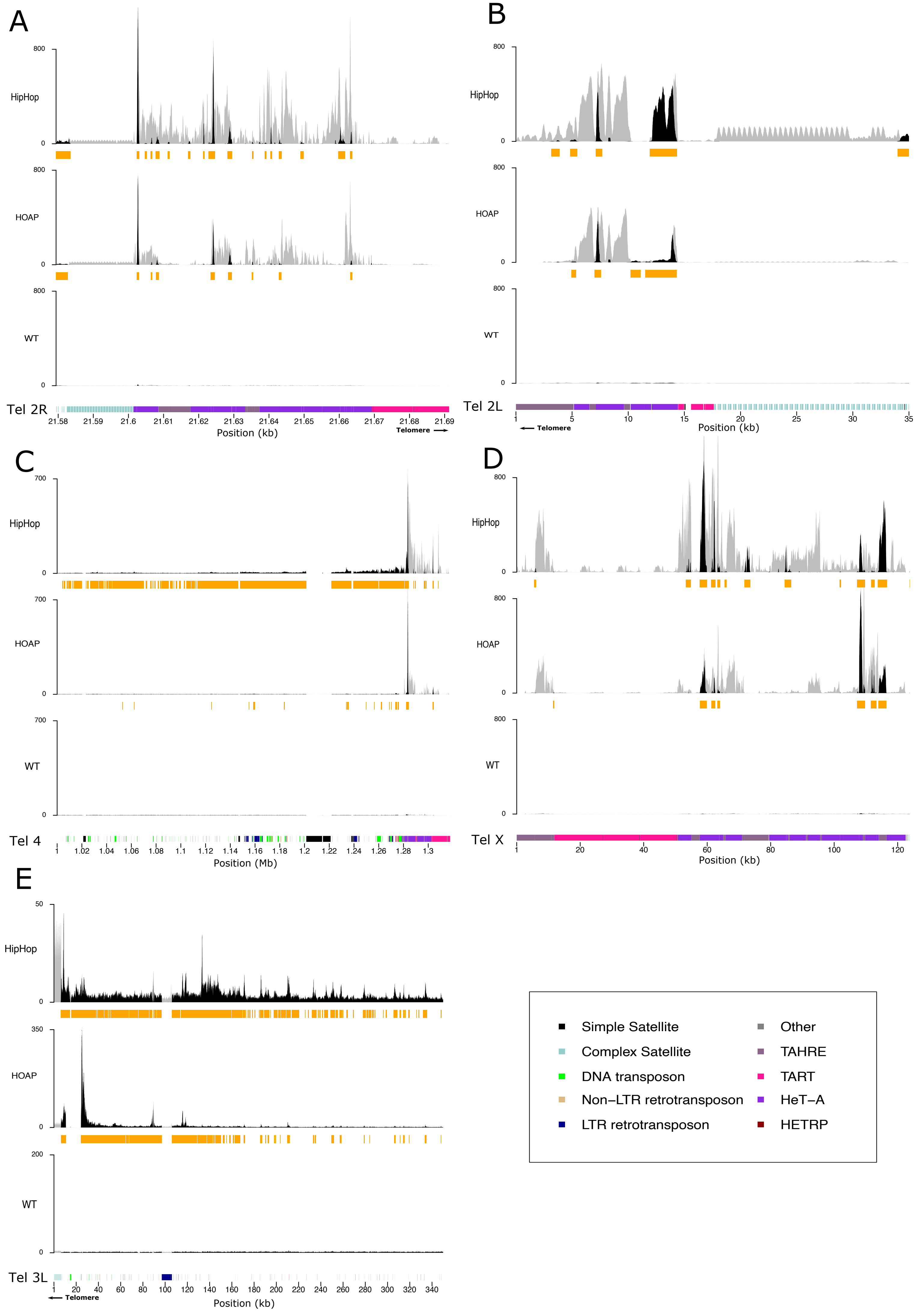

### supplemental figure S4

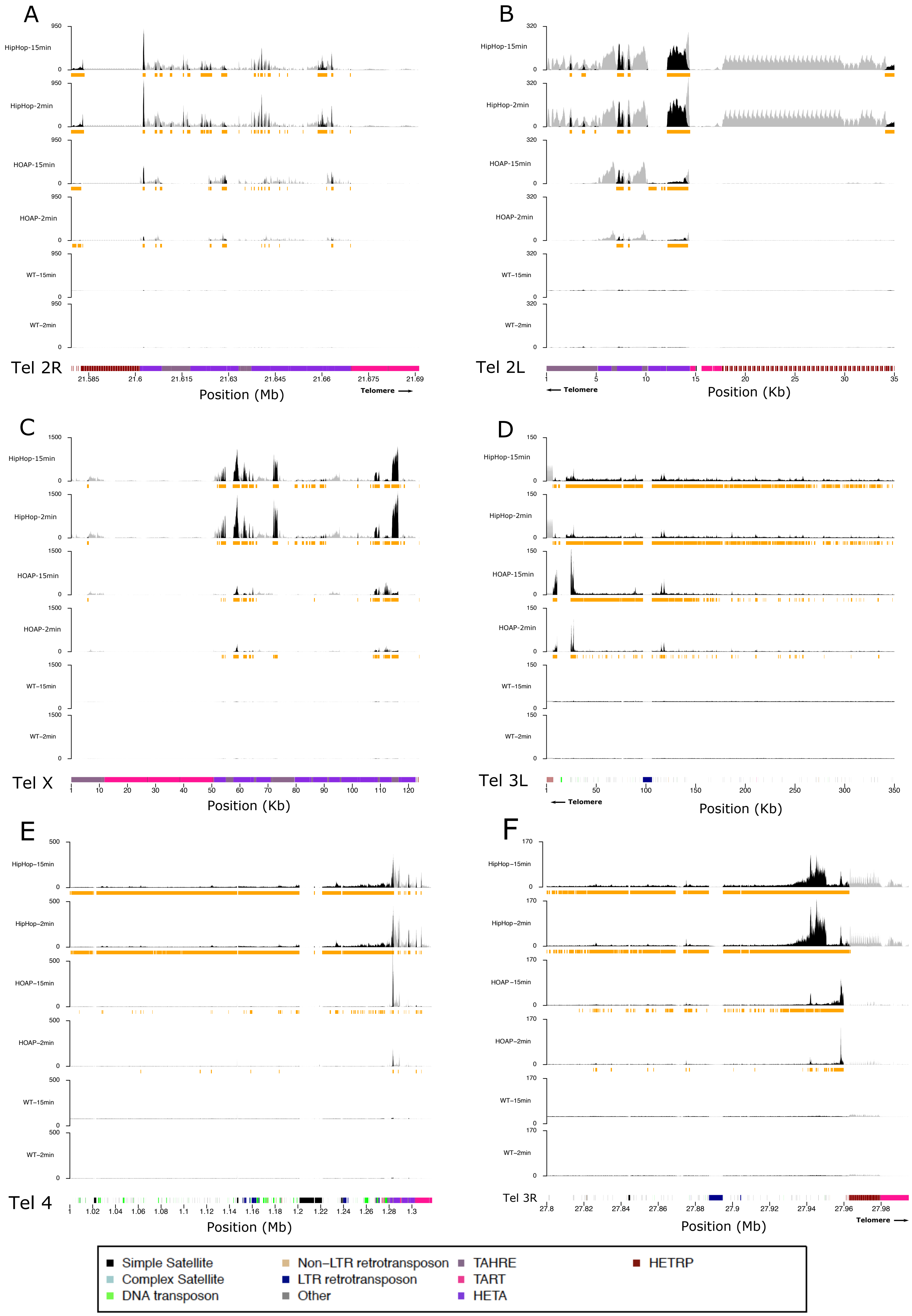

### supplemental figure S5

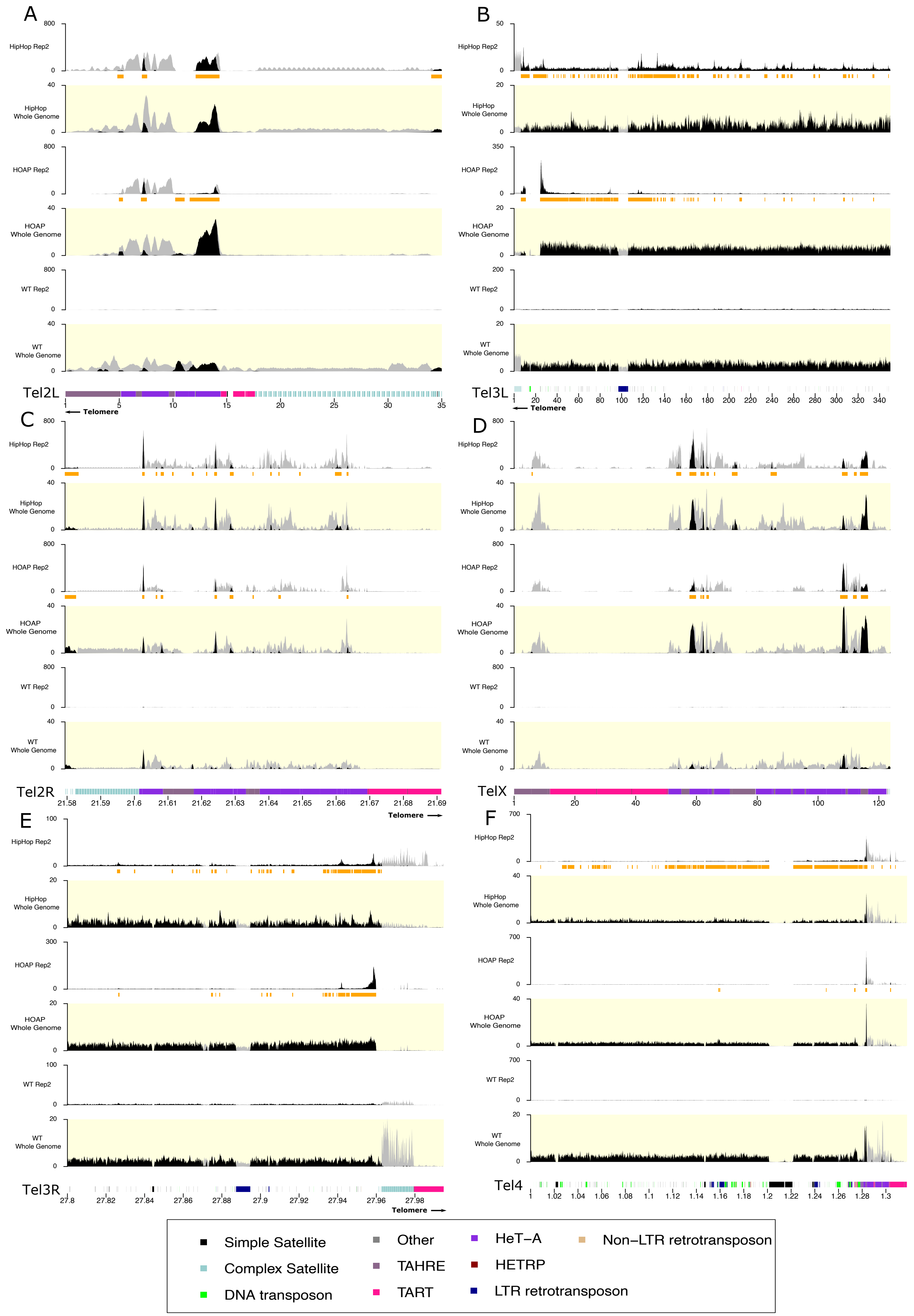

### supplemental figure S6

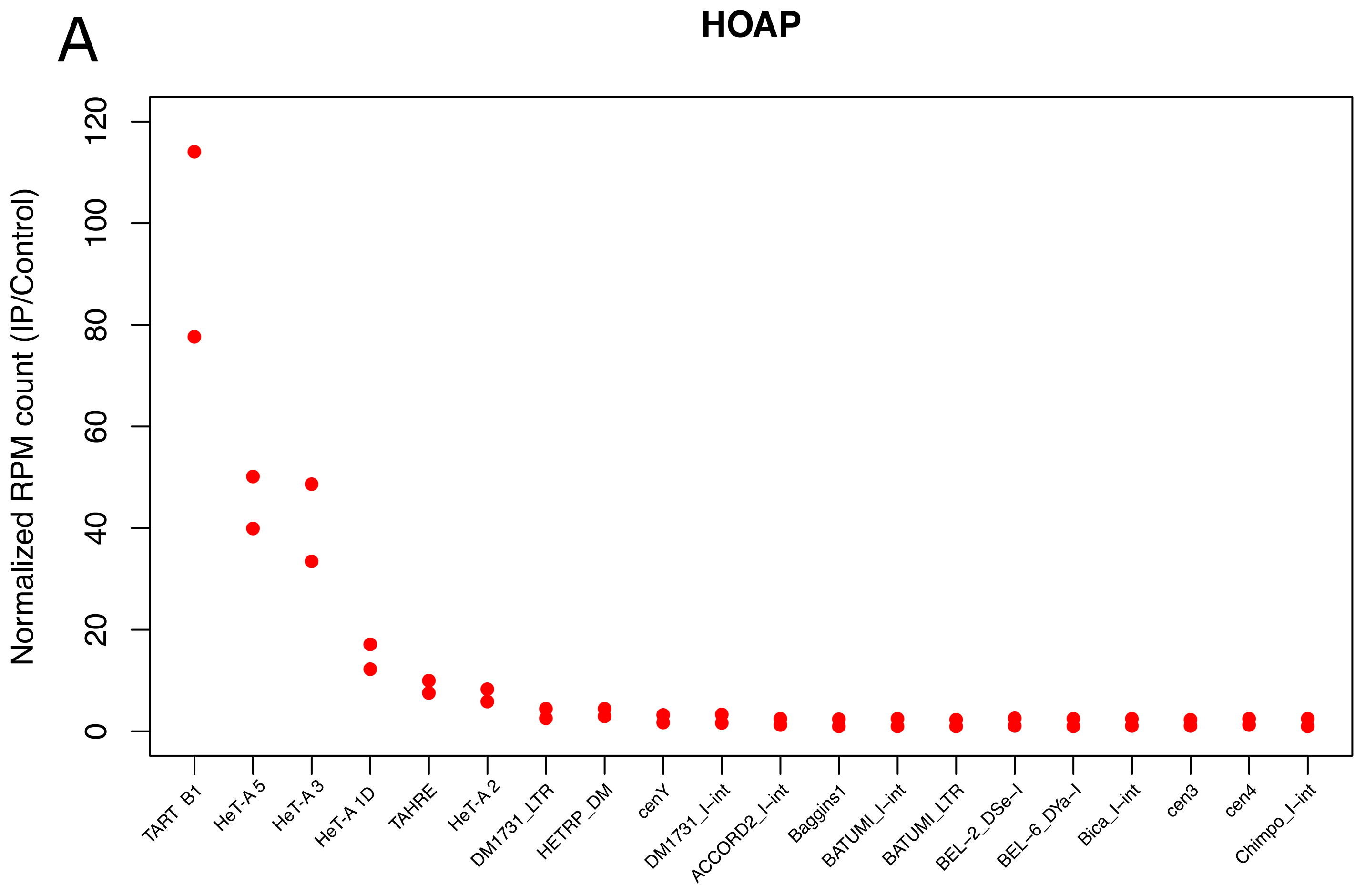

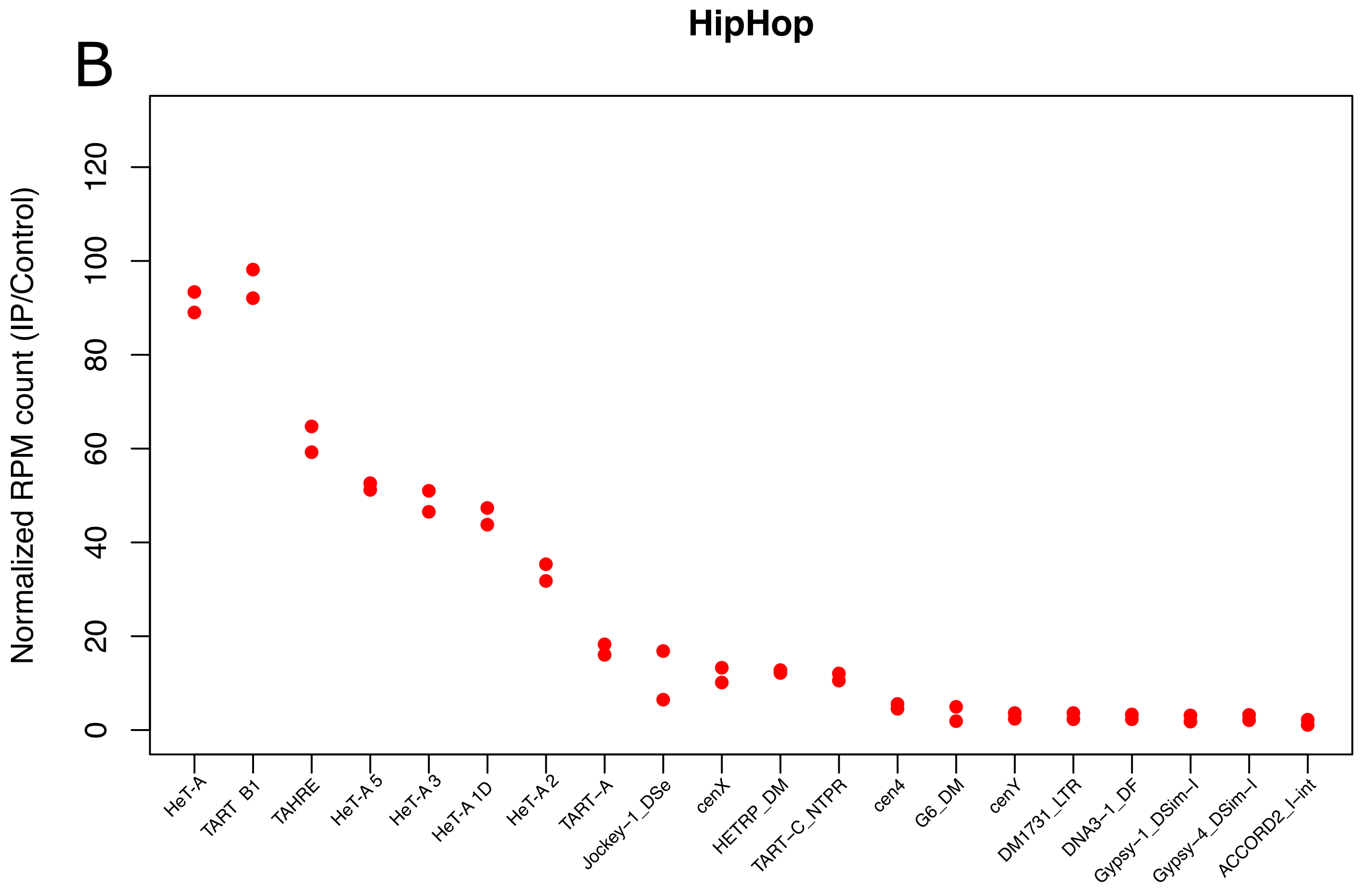

### supplemental figure S7

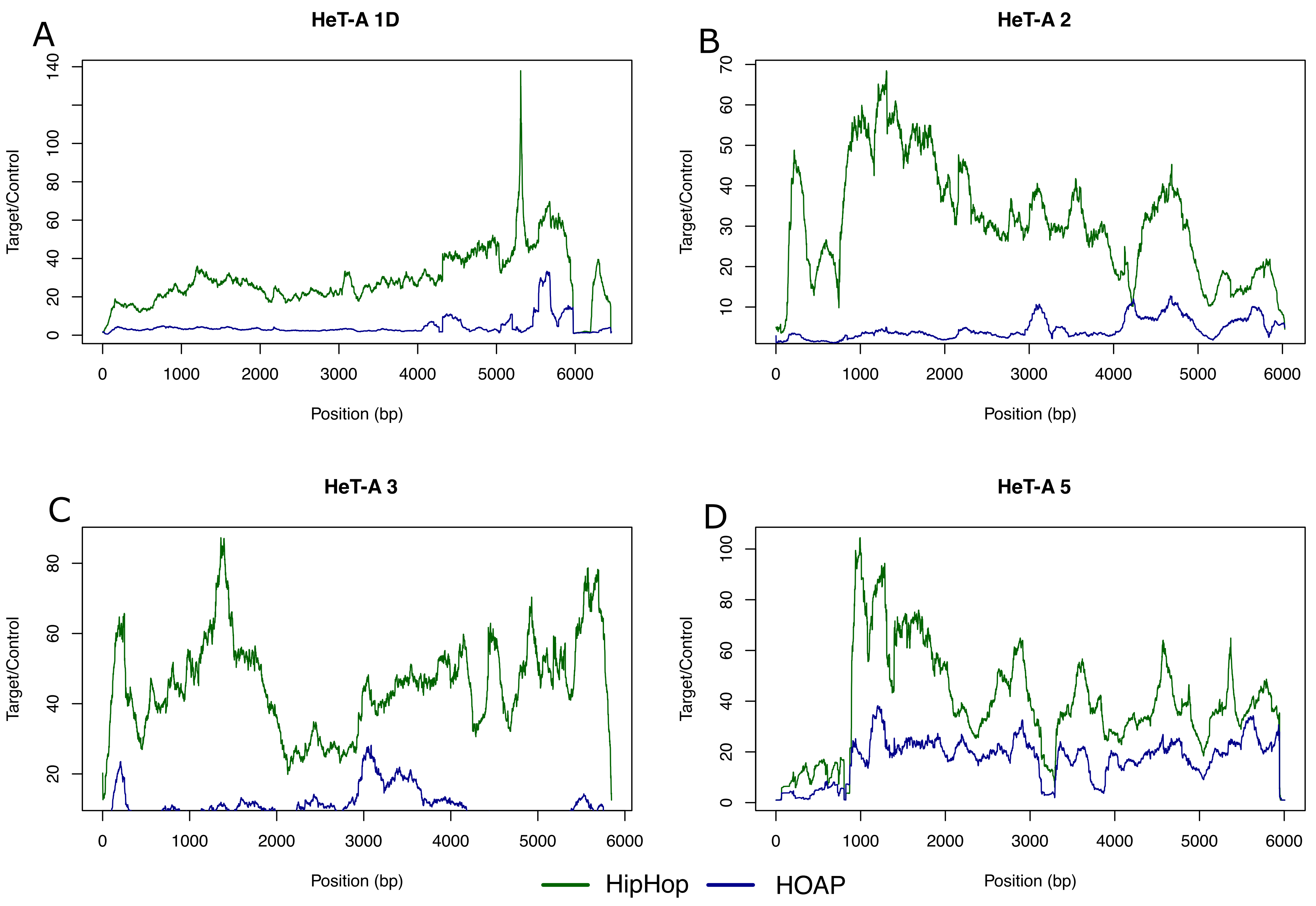

### supplemental figure S8

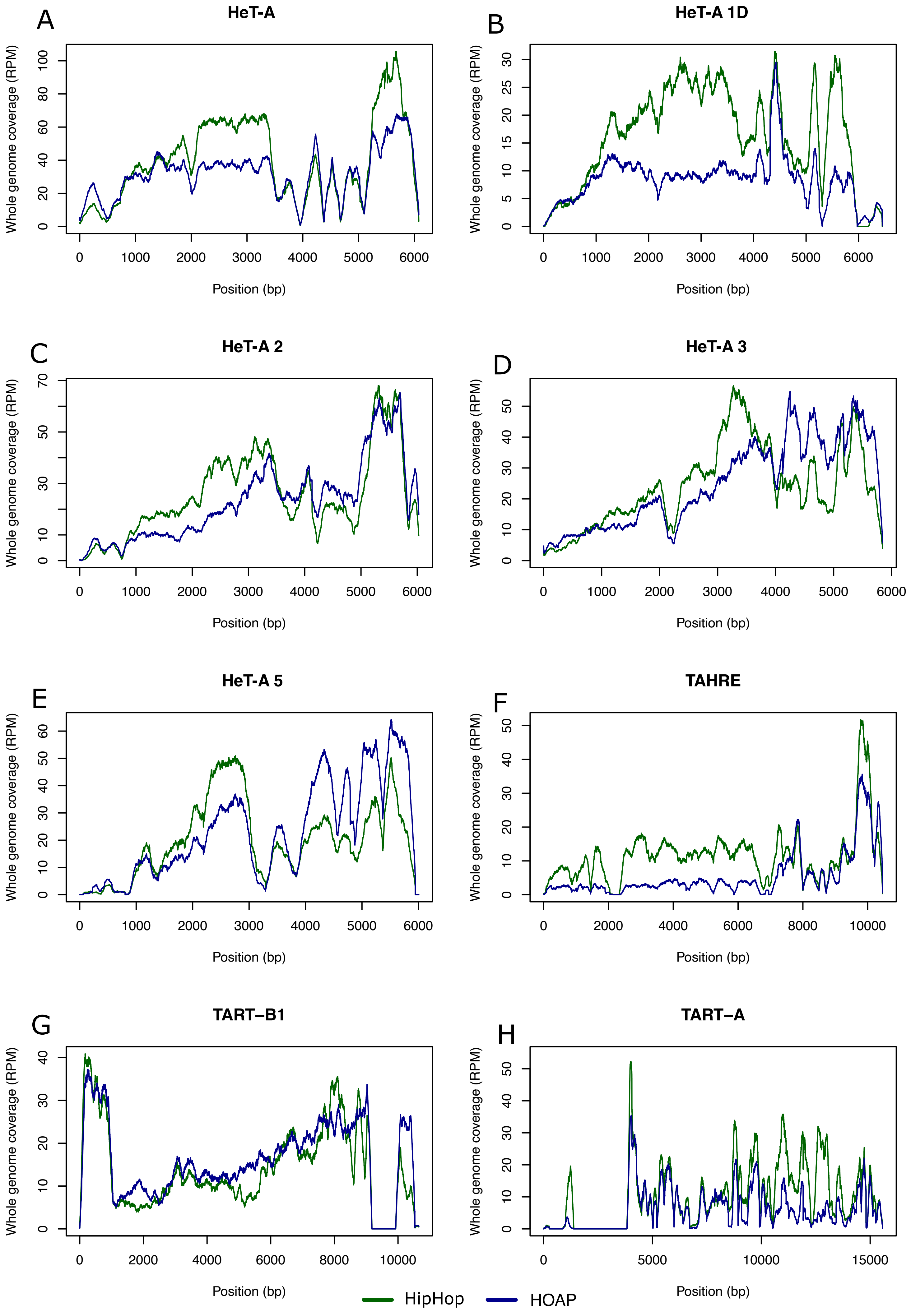

### supplemental figure S9

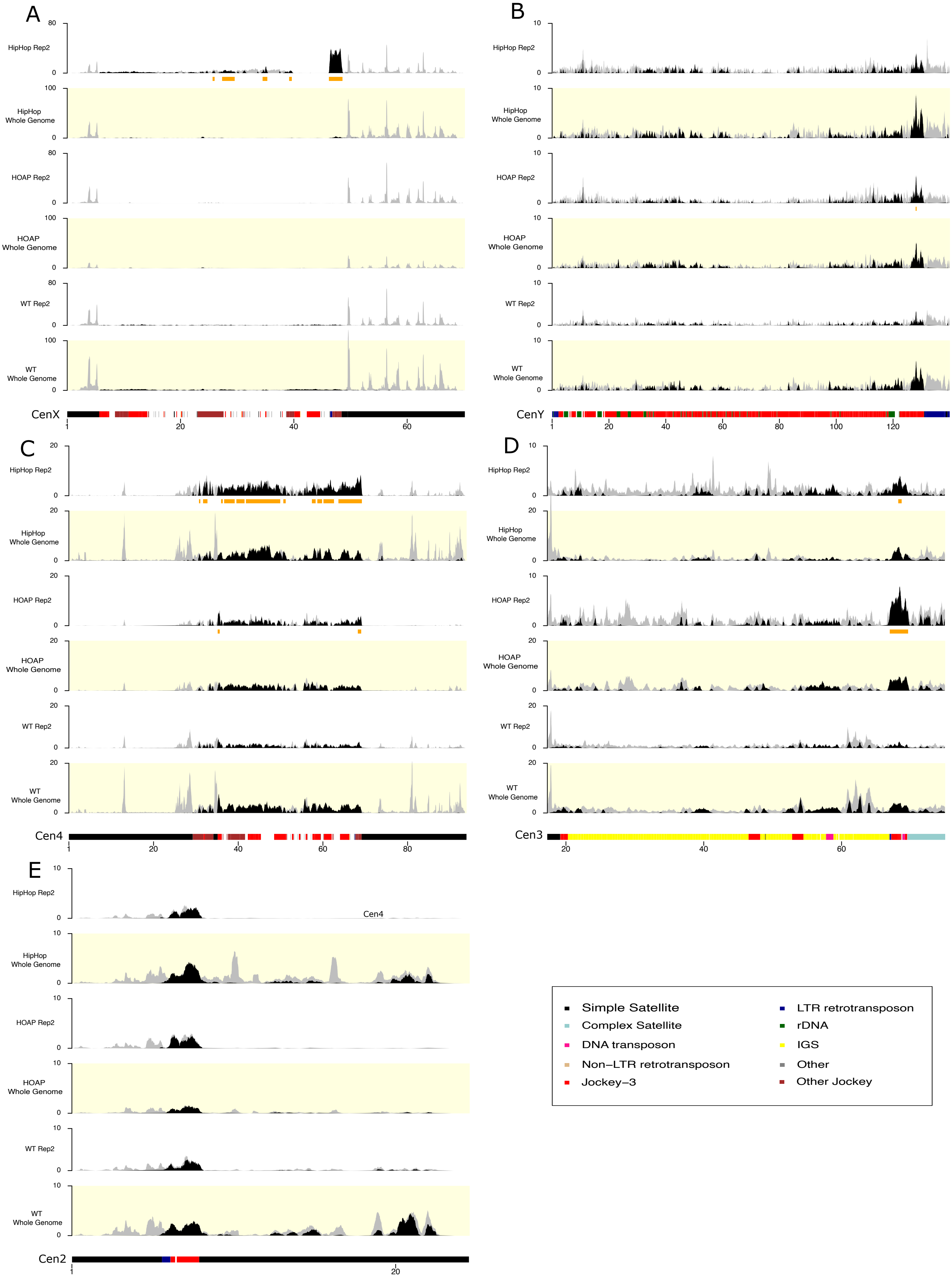
